# Nucleosome-CHD4 chromatin remodeller structure explains human disease mutations

**DOI:** 10.1101/665562

**Authors:** Lucas Farnung, Moritz Ochmann, Patrick Cramer

## Abstract

Chromatin remodelling plays important roles in gene regulation during development, differentiation and in disease. The chromatin remodelling enzyme CHD4 is a component of the NuRD and ChAHP complexes that are involved in gene repression. Here we report the cryo-electron microscopy (cryo-EM) structure of *Homo sapiens* CHD4 engaged with a nucleosome core particle in the presence of the non-hydrolysable ATP analogue AMP-PNP at an overall resolution of 3.1 Å. The ATPase motor of CHD4 binds and distorts nucleosomal DNA at super-helical location (SHL) +2, supporting the ‘twist defect’ model of chromatin remodelling. CHD4 does not induce unwrapping of terminal DNA, in contrast to its homologue Chd1, which functions in gene activation. Our results also rationalize the effect of CHD4 mutations that are associated with cancer or the intellectual disability disorder Sifrim-Hitz-Weiss syndrome.

## Introduction

In the nucleus of eukaryotic cells, DNA is compacted into chromatin. The fundamental building block of chromatin is the nucleosome, a complex of 146 base pairs (bp) of DNA wrapped around an octamer of histone proteins. The degree of chromatin compaction influences DNA replication, transcription, and repair. Maintenance of the appropriate chromatin state requires ATP-dependent chromatin-remodelling enzymes. These ‘chromatin remodellers’ are divided into four families, called CHD, SWI/SNF, ISWI, and INO80 (Clapier et al., 2017). All chromatin remodellers contain a conserved ATPase core that utilizes ATP hydrolysis to alter contacts between nucleosomal DNA and the histone octamer and to facilitate nucleosome assembly, sliding, ejection, or histone exchange.

Members of the CHD (chromodomain helicase DNA-binding) protein family of chromatin remodellers all contain a central SNF2-like ATPase motor domain and a double chromodomain in their N-terminal region. The double chromodomain binds modified histones(Sims et al., 2005) and interacts with nucleosomal DNA (Nodelman et al., 2017). The interaction with DNA regulates and fine tunes ATPase activity. Recent structures of the yeast remodeller Chd1 in complex with a nucleosome uncovered the architecture of one subfamily of CHD remodellers (subfamily I) and its interactions with the nucleosome (Farnung et al., 2017; Sundaramoorthy et al., 2018). A unique feature of these structures is that Chd1 binding induces unwrapping of terminal DNA from the histone octamer surface at superhelical location (SHL) -6 and -7 (Farnung et al., 2017; Sundaramoorthy et al., 2018). However, the resolution of these studies was limited, such that atomic details were not resolved.

The human CHD family member CHD4 (Woodage et al., 1997) shows nucleosome spacing activity (Silva et al., 2016). CHD4 is also known as Mi-2 in *Drosophila melanogaster* (Kehle et al., 1998) and together with CHD3 forms subfamily II, which differs in domain architecture from subfamily I. CHD3 and CHD4 contain two N-terminal plant homeodomain zinc fingers (Schindler et al., 1993) (PHD fingers 1 and 2), a DNA-interacting double chromodomain, and the ATPase motor. CHD4 contains an additional high mobility group (HMG) box-like domain in its N-terminal region (Silva et al., 2016) and two additional domains of unknown function located in the C-terminal region.

CHD4 is implicated in the repression of lineage-specific genes during differentiation (Liang et al., 2017) and is required for the establishment and maintenance of more compacted chromatin structures (Bornelöv et al., 2018). CHD4 mutations have a high incidence in some carcinomas (Getz et al., 2013) as well as thyroid and ovarian cancers (Längst and Manelyte, 2015). Some mutations in CHD4 have also been implicated in intellectual disability syndromes (Sifrim et al., 2016; Weiss et al., 2016). CHD4 is part of the multi-subunit Nucleosome Remodelling Deacetylase (NuRD) complex (Tong et al., 1998; Xue et al., 1998; Zhang et al., 1998). NuRD also contains the deacetylase HDAC1/2 and accessory subunits that serve regulation and scaffolding roles. NuRD is implicated in gene silencing, but also gene activation (Gnanapragasam et al., 2011). It is essential for cell cycle progression (Polo et al., 2010), DNA damage response (Larsen et al., 2010; Smeenk et al., 2010), establishment of heterochromatin (Sims and Wade, 2011), and differentiation (Bornelöv et al., 2018; Burgold et al., 2019). It was recently shown that CHD4 is also part of the heterotrimeric ChAHP complex that is also involved in gene repression (Ostapcuk et al., 2018).

Thus far, structural studies of CHD4 have been limited to individual domains (Kwan et al., 2003; Mansfield et al., 2011). Here we report the cryo-electron microscopy (cryo-EM) structure of human CHD4 bound to a nucleosome at an overall resolution of 3.1 Å. CHD4 engages the nucleosome at SHL +2 and induces a conformational change in DNA at this location in the presence of the ATP analogue adenylyl imidodiphosphate (AMP-PNP). Structural comparisons show that CHD4, in contrast to Chd1, does not induce unwrapping of terminal DNA. Maintenance of the integrity of the nucleosome in the presence of CHD4 is consistent with the role of CHD4 in gene repression, and in heterochromatin formation and maintenance. Finally, the detailed nucleosome-CHD4 structure enables mapping of known human disease mutations (Kovač et al., 2018; Sifrim et al., 2016; Weiss et al., 2016) and indicates how these perturb enzyme function.

## Results

### Nucleosome-CHD4 complex structure

To investigate how the human chromatin remodeller CHD4 engages a nucleosome and to understand the structural basis of cancer-related mutations in CHD4, we determined the structure of *H. sapiens* CHD4 bound to a Xenopus laevis nucleosome core particle in the presence of the ATP analogue AMP-PNP. We recombinantly expressed and purified full-length CHD4 and reconstituted a complex of CHD4 with a pre-assembled nucleosome core particle. The nucleosome comprised 145 base pairs (bp) of DNA, corresponding to the Widom 601 sequence (Lowary and Widom, 1998) with additional 4 and 30 bp of extranucleosomal DNA on the entry and exit side of the nucleosome, respectively. The nucleosome-CHD4 complex was purified by size exclusion chromatography (Supplementary Fig. 1).

To determine the structure of the nucleosome-CHD4 complex, we collected single particle cryo-EM data on a Titan Krios (FEI) microscope equipped with a K2 direct electron detector (Gatan) (Methods). We obtained a cryo-EM reconstruction of the nucleosome-CHD4 complex at an overall resolution of 3.1 Å (FSC 0.143 criterion) (Supplementary Fig. 2-4). The nucleosome was resolved at a resolution of 3.0-4.5 Å, whereas CHD4 was resolved at 3.1-5.0 Å depending on the protein region. The register of the DNA was unambiguously determined based on distinct densities for purine and pyrimidine nucleotides around the dyad axis (Supplementary Fig. 3h). Well-defined density was also obtained for AMP-PNP and a coordinated magnesium ion in the CHD4 active site (Supplementary Fig. 3i). The model was locally adjusted and real-space refined, leading to very good stereochemistry (Methods) (Table 1).

**Table 1.** Cryo-EM data collection, refinement and validation statistics.

**Tables and Supplementary Figures**
|  | Nucleosome-CHD4<br>complex<br>(EMD-10058)<br>(PDB 6RYR) | Nucleosome-CHD4<sub>2</sub> complex<br>(EMDB-10059)<br>(PDB 6RYU) |
| --- | --- | --- |
| **Data collection and processing** |  |  |
| Magnification | 130,000 | 130,000 |
| Voltage (kV) | 300 | 300 |
| Electron exposure (e<sup>-</sup>/Å<sup>2</sup>) | 43-45 | 43-45 |
| Defocus range (μm) | 0.25-4 | 0.25-4 |
| Pixel size (Å) | 1.05 | 1.05 |
| Symmetry imposed | C1 | C1 |
| Initial particle images (no.) | 650,599 | 650,599 |
| Final particle images (no.) | 89,623 | 40,233 |
| Map resolution (Å) | 3.1 | 4.0 |
| FSC threshold | 0.143 | 0.143 |
| Map resolution range (Å) | 3.0-5 | 3.7-8.3 |
| **Refinement** |  |  |
| Initial models used (PDB code) | 3LZ0, 5O9G, 2L75, 4O9I | 3LZ0, 5O9G, 2L75, 4O9I |
| Map sharpening *B* factor (Å<sup>2</sup>) | -36 | -86 |
| Model composition |  |  |
| Non-hydrogen atoms | 17,837 | 23,614 |
| Protein residues | 1463 | 2168 |
| Nucleotides | 298 | 298 |
| Ligands | 4 | 8 |
| *B* factors (Å<sup>2</sup>) |  |  |
| Protein | 33.37 | 79.70 |
| Nucleotide | 70.91 | 104.76 |
| Ligand | 44.86 | 117.67 |
| R.m.s. deviations |  |  |
| Bond lengths (Å) | 0.008 | 0.007 |
| Bond angles (°) | 0.910 | 0.958 |
| **Validation** |  |  |
| MolProbity score | 1.59 | 1.90 |
| Clashscore | 4.08 | 7.80 |
| Poor rotamers (%) | 0.16 | 0.66 |
| Ramachandran plot |  |  |
| Favored (%) | 94.15 | 92.31 |
| Allowed (%) | 5.71 | 7.64 |
| Disallowed (%) | 0.14 | 0.05 |

### CHD4 architecture

The CHD4 ATPase motor binds the nucleosome at SHL +2 (Fig. 1). Binding at this location has also been observed for the chromatin remodellers Chd1 (Farnung et al., 2017; Sundaramoorthy et al., 2018), Snf2 (Liu et al., 2017), and Swr1 (Willhoft et al., 2018). The ATPase motor is in a closed, post-translocated state with AMP-PNP bound in the active site. A similar state was observed for Chd1 when bound to ADP!·BeF_3_ (Farnung et al., 2017; Sundaramoorthy et al., 2018, 2017). The double chromodomain is located at SHL +1 and contacts the nucleosomal DNA phosphate backbone via electrostatic interactions, in a fashion similar to that observed for *S. cerevisiae* Chd1 (Fig. 1) (Farnung et al., 2017; Nodelman et al., 2017).

**Fig. 1.**
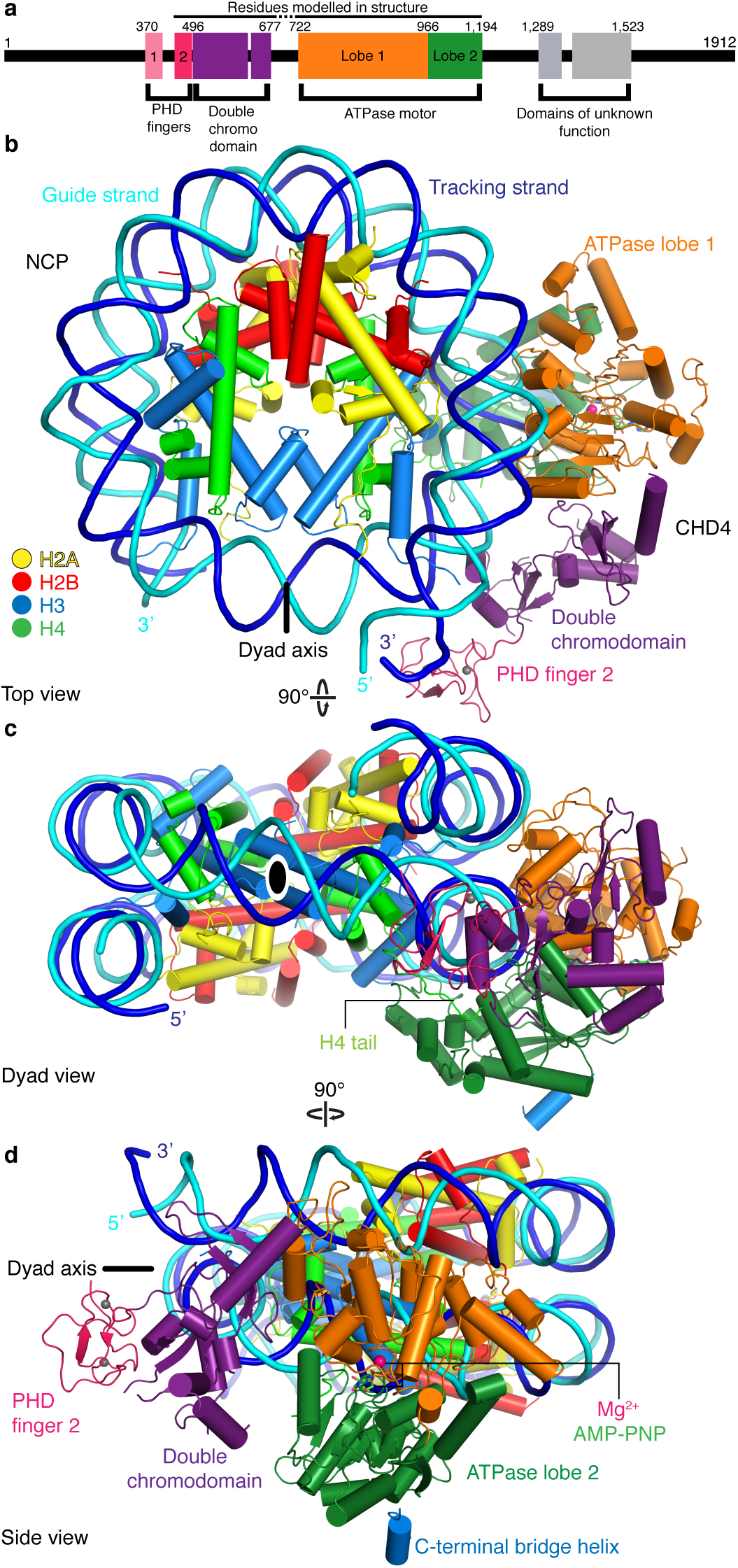
Structure of the nucleosome-CHD4 complex. a, Schematic of domain architecture of CHD4. Domain borders are indicated. b-d, Cartoon model viewed from the top (b), dyad (c), and side (d). Histones H2A, H2B, H3, H4, tracking strand, guide strand, CHD4 PHD finger 2, double chromodomain, ATPase lobe 1, and ATPase lobe 2 are coloured in yellow, red, light blue, green, blue, cyan, pink, purple, orange, and forest green, respectively. Colour code used throughout. The dyad axis is indicated as a black line or a black oval circle. Magnesium and zinc ions shown as pink and grey spheres, respectively. AMP-PNP shown in stick representation.

The CHD4 domain PHD finger 2 is located near SHL +0.5 and the double chromodomain. This location is consistent with NMR studies that predicted binding of this PHD finger close to the dyad axis and the H3 tail (Gatchalian et al., 2017). Additionally, we observe parts of the C-terminal bridge (Hauk et al., 2010), an amino acid segment that follows the ATPase lobe. Part of the C-terminal bridge docks against ATPase lobe 2 and extends towards the first ATPase lobe (Fig. 1, Supplementary Figure 3j). This region was not resolved in the nucleosome-Chd1 structures but was observed in a previously published crystal structure of auto-inhibited Chd1 (Hauk et al., 2010). Taken together, CHD4 and Chd1 share a core architecture that involves the ATPase motor and the double chromodomain but differ in their peripheral subfamily-specific protein features.

### CHD4 binding does not detach exit side nucleosomal DNA

In contrast to the nucleosome-Chd1 structure (Farnung et al., 2017), we did not observe unwrapping of nucleosomal DNA from the histone octamer on the second DNA gyre at SHL -6 and -7 (Fig. 2). This major difference between these complex structures may be due to a lack of a DNA-binding region in CHD4. Chd1 uses its DNA-binding region to interact extensively with terminal DNA on the exit side at SHL -7, and such contacts are absent in the nucleosome-CHD4 structure (Fig. 2). It is likely that other CHD family members such as CHD3 and CHD5, which also lack a DNA-binding region, will also not induce unwrapping of terminal DNA.

**Fig. 2.**
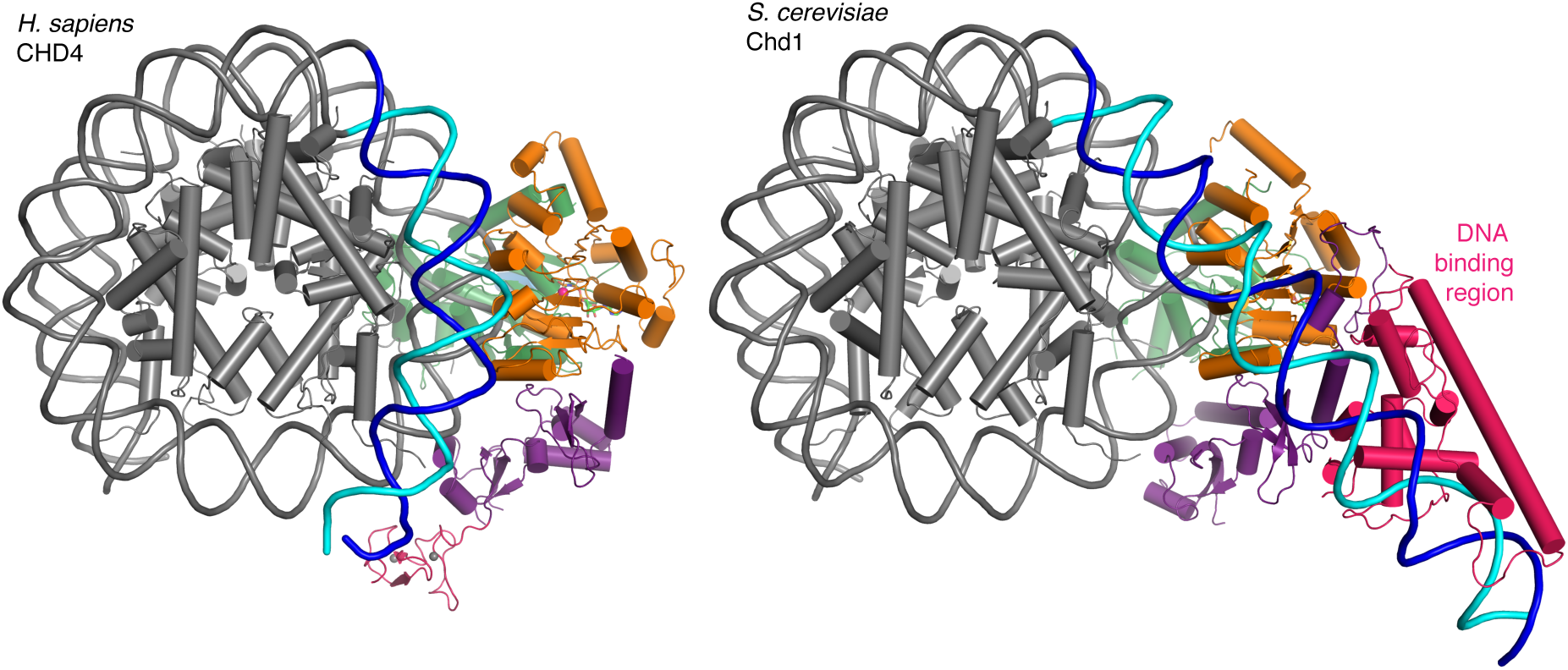
Comparison with nucleosome-Chd1 structure. CHD4 (left) does not possess a DNA-binding region and does not detach DNA from the second gyre. Chd1 (right) detaches DNA from SHL -7 to -5, stabilizes the detached DNA via its DNA binding region, and introduces a ∼60°bend with respect to the canonical DNA position observed in the nucleosome-CHD4 structure.

### CHD4-DNA interactions

The high resolution of our nucleosome-CHD4 structure enables a detailed description of the interactions of the ATPase motor with nucleosomal DNA. CHD4 contacts the phosphate backbone of the tracking and guide strand via electrostatic interactions that are mostly mediated by lysine and arginine residues (Fig. 3). These interactions with the DNA phosphate backbone are formed by residues in the canonical ATPase motifs Ia, Ic, II, IV, IVa, V, and Va and by residues present in non-canonical motifs (e.g. Lys810) (Fig. 3, Supplementary Fig. 5).

**Fig. 3.**
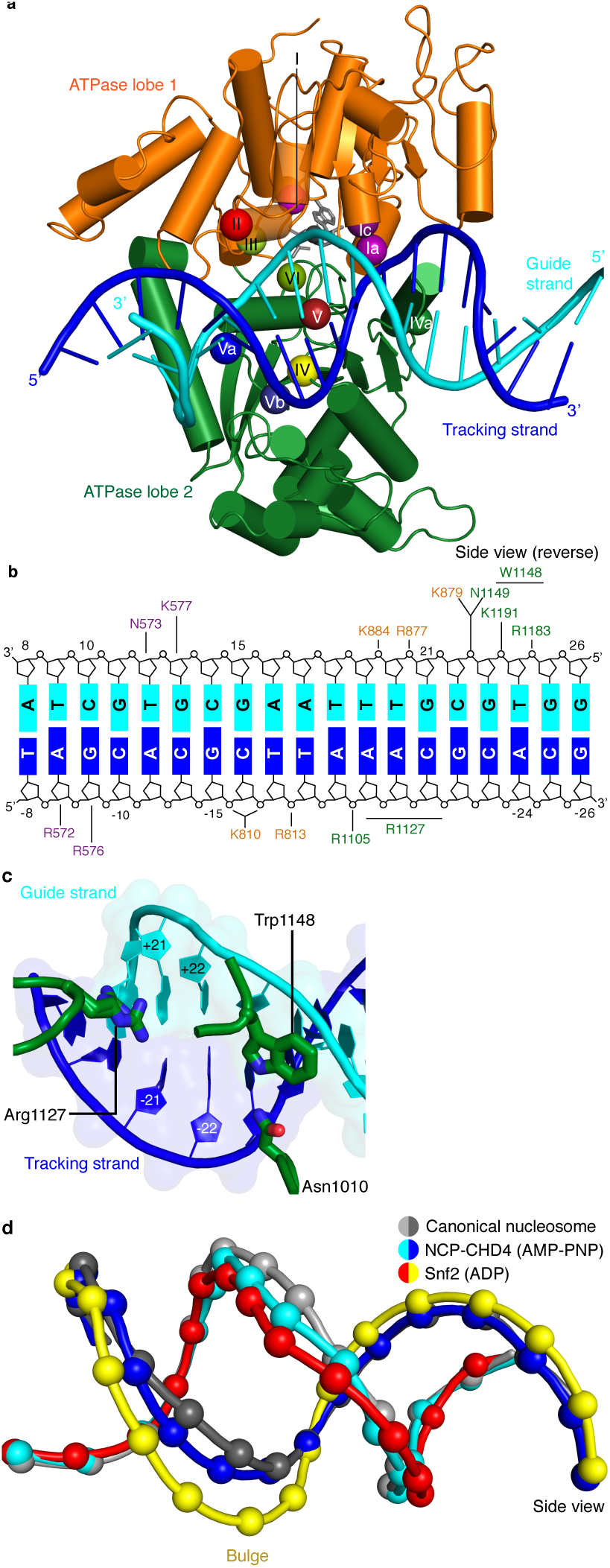
CHD4-DNA interactions and DNA distortion. a, CHD4 interacts extensively with nucleosomal DNA around SHL +2. ATPase lobe 1 and lobe 2 of CHD4 are shown. Guide and tracking strands are indicated. ATPase motifs are shown as coloured spheres and labelled. b, Schematic depiction of DNA interactions of the double chromodomain, ATPase lobe 1 and lobe 2. c, Asn1010, Trp1148 and Arg1227 insert into the minor groove between DNA tracking and guide strand. The two conformations of the Arg1127 side chain are shown. Nucleic acids are shown as cartoons with their respective surfaces. d, Detailed cartoon representation of DNA distortion at SHL +2. Canonical nucleosome (PDB code 3LZ0, grey), AMP-PNP bound NCP-CHD4 structure (this study, blue and cyan), and ADP bound nucleosome-Snf2 structure (PDB code 5Z3O, red and yellow) are shown. Phosphate atoms shown as spheres.

We also observe that residues Asn1010, Arg1127, and Trp1148 insert into the DNA minor groove over a stretch of seven base pairs (Fig. 3c). Asn1010 is not part of a canonical ATPase motif and inserts into the DNA minor groove around SHL +2.5. Arg1127 (motif V) is universally conserved in all CHD chromatin remodellers and inserts into the DNA minor groove at SHL +2. Our density is consistent with two alternative conformations of the Arg1127 side chain, with the guanidinium head group pointing either towards the tracking or the guide strand of DNA. Trp1148 is located in motif Va, inserts into the minor groove near the guide strand, and plays a critical role in coupling ATPase hydrolysis and DNA translocation (Liu et al., 2017). We further observe a contact between a negatively charged loop in ATPase lobe 1 (residues 832-837) and the second DNA gyre at SHL -6. This loop is present in CHD3, CHD4, and CHD5, but not in Snf2 or ISWI remodellers (Supplementary Fig. 5).

### CHD4 binding distorts DNA at SHL +2

Comparison of our structure with a high-resolution X-ray crystallographic structure of the free nucleosome (Vasudevan et al., 2010) reveals a conformational change in the DNA where the ATPase motor engages its DNA substrate (SHL +2) (Fig. 3d). The high resolution of the nucleosome-CHD4 structure shows that 5 DNA base pairs between SHL +1.5 and SHL +2.5 are pulled away from the octamer surface by up to 3 Å. This distortion does not include the previously observed ‘bulging’ or a ‘twist defect’ that is characterized by a 1 bp local underwinding of the DNA duplex and observed when the ATPase motor adopts the open/apo or ADP-bound states (Li et al., 2019). In contrast, the DNA distortion observed in our AMP-PNP bound state is an intermediate between the bulged and the canonical DNA conformation (Fig. 3d). This AMP-PNP bound intermediate DNA state was predicted based on biochemical experiments (Winger et al., 2018). This observation demonstrates that the extent of DNA distortion at SHL +2 depends on the functional state of the ATPase motor and is consistent with the proposed twist defect propagation model of chromatin remodelling (Winger et al., 2018).

### CHD4 binds the histone H4 tail

As observed for *S. cerevisiae* Chd1 (Farnung et al., 2017), *H. sapiens* CHD4 contacts the histone H4 tail with its ATPase lobe 2. The H4 tail is located between ATPase lobe 2 and the nucleosomal DNA at SHL +1.5. The conformation of the H4 tail differs from that observed in structures of the free nucleosome where the tail makes inter-nucleosomal contacts with the ‘acidic patch’ of a neighbouring nucleosome. It also differs from the H4 position observed in a higher-order structure where the H4 tail extends over the DNA interface between two nucleosomes (Schalch et al., 2005). A loop in lobe 2 of the ATPase (CHD4 residues 1001-1006) replaces the H4 tail in this position, apparently inducing H4 positioning that allows ATPase lobe 2 binding (Fig. 4a).

**Fig. 4.**
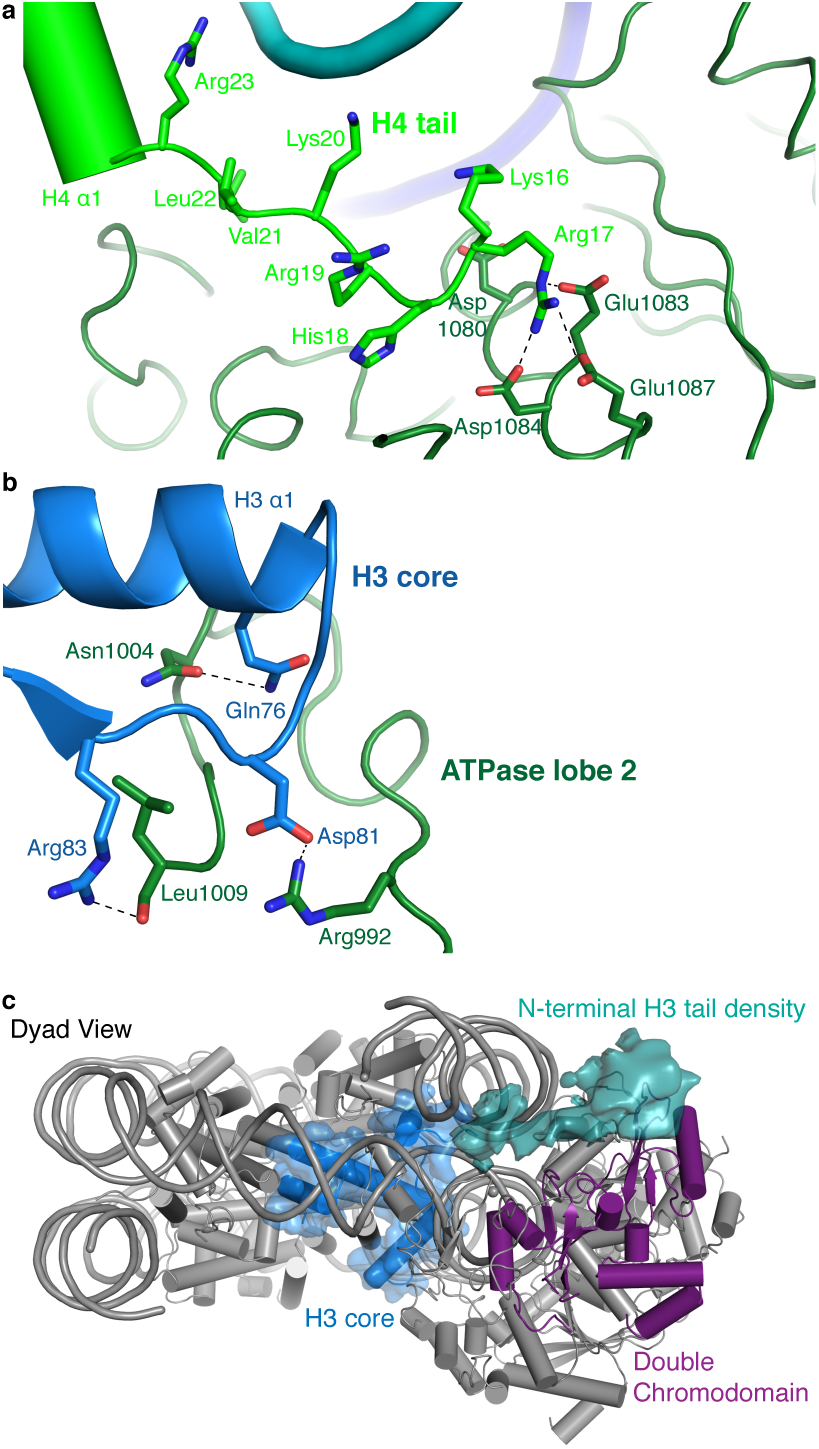
CHD4 contacts H3 and H4. a, ATPase lobe 2 interacts extensively with the H4 tail. b, A loop in ATPase lobe 2 contacts H3 alpha helix 1 and neighbouring residues. c, The double chromodomain of CHD4 contacts the H3 N-terminal tail. H3 core is shown in blue, H3 tail density from the low-pass filtered final map (7 Å) in teal, and the double chromodomain in purple.

ATPase lobe 2 contains a highly acidic cavity formed by Asp1080, Glu1083, Asp1084, and Glu1087 (Fig. 4a). This acidic cavity is conserved across all CHD family members. The basic side chain of the H4 histone tail residue Arg17 inserts into this acidic cavity (Fig. 4a). Similar interactions with the H4 tail have also been reported for Snf2 and ISWI remodellers (Armache et al., 2019; Yan et al., 2019). The side chain of H4 Lys16 also points towards the acidic cavity and is positioned in close proximity to residues Asp1080 and Glu1083. Acetylation of H4 Lys16 is therefore predicted to weaken these charge-based interactions and to reduce the affinity of chromatin remodellers for the H4 tail, as noted before (Yan et al., 2016).

### CHD4 interacts with histone H3

The ATPase lobe 2 also contacts the core of histone H3 (alpha helix 1, Gln76 and Arg83) via CHD4 residues Asn1004 and Leu1009, respectively (Fig. 4b). This contact is critical for chromatin remodelling. Deletion of the homologous region in Chd1 leads to abolishment of chromatin remodelling activity (Sundaramoorthy et al., 2018). However, it remains unclear if these contacts are required for proper substrate recognition and positioning or whether they are also necessary to generate the force required for DNA translocation. Low-pass filtering of our map further shows the H3 N-terminal tail trajectory, which extends to the double chromodomain (Fig. 4c). The contact between the H3 tail and the double chromodomain could target CHD4 to nucleosomes methylated at Lys27 of H3 (Kuzmichev et al., 2002), a classical mark for gene repression.

### Two CHD4 molecules can engage with the nucleosome

During 3D classification of our cryo-EM dataset we observed a distinct class of particles that showed two CHD4 molecules bound to the same nucleosome (Fig. 5, Supplementary Fig. 2-4). Refinement of this class of particles yielded a reconstruction at an overall resolution of 4.0 Å (FSC 0.143 criterion) (Table 1). A model of this nucleosome-CHD42 complex was obtained by docking the refined nucleosome-CHD4 model into the density and then placing another CHD4 molecule into the additional density observed on the opposite side. The resulting nucleosome-CHD42 complex structure shows pseudo-twofold symmetry with CHD4 molecules bound at SHL +2 and SHL -2 (Fig. 5). The second CHD4 molecule uses its double chromodomain and PHD finger 2 to contact nucleosomal DNA at SHL +1 and +0.5, respectively. Binding of the second CHD4 molecule also did not lead to unwrapping of terminal DNA. Binding of two chromatin remodellers to a single nucleosome was previously observed for *S. cerevisiae* Chd1 (Sundaramoorthy et al., 2018) and *H. sapiens* SNF2H (Armache et al., 2019). However, in contrast to the structure of the nucleosome-SNF2H2 complex, we do not observe a distortion in the histone octamer due to the presence of the chromatin remodellers. Binding of two remodeller molecules could allow for higher efficiency in positioning the nucleosome at a precise location but necessitates coordination of the remodellers. A possible mechanism for coordination could be that twist defects that are introduced by remodeller binding are propagated from the entry SHL 2 into the exit side SHL 2 (Brandani et al., 2018; Brandani and Takada, 2018). Presence of the twist defect at the second remodeller binding site could interfere with the translocation activity of the second remodeller (Sabantsev et al., 2019).

**Fig. 5.**
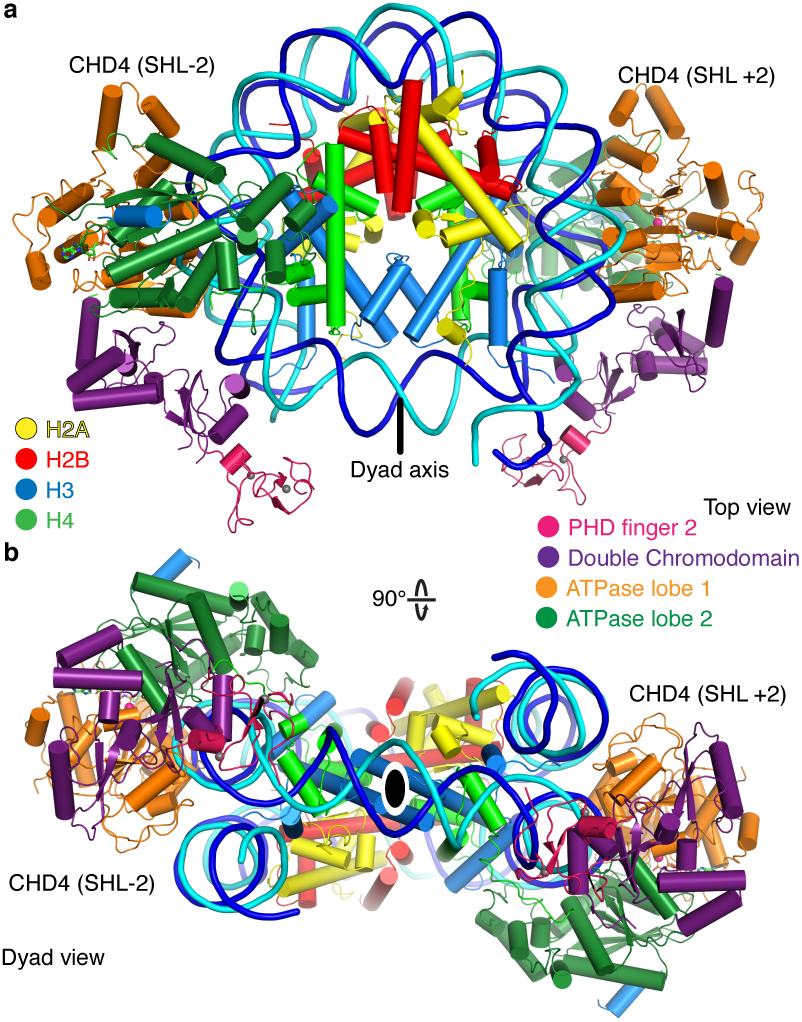
The nucleosome can bind two copies of CHD4. Cartoon model of the nucleosome-CHD4_2_ structure viewed from the top (a), and dyad view (b).

### Cancer-related CHD4 mutations

Many studies have reported mutations in CHD4 that are related to human diseases, in particular cancer (Xia et al., 2017). Mutations involved in various cancer phenotypes have been observed in the PHD finger 2, the double chromodomain, and both lobes of the ATPase motor. To elucidate effects of such mutations on CHD4 activity, the *D. melanogaster* CHD4 homologue Mi-2 has been used as a model protein for functional analysis (Kovač et al., 2018). CHD4 mutations have been found to fall in two categories. Whereas some mutations influence ATPase and DNA translocation activity (Arg1162, His1196, His1151 and Leu1215), other mutations seem to change protein stability (Leu912, and Cys464) or disrupt DNA binding (Val558 and Arg572).

To rationalize these findings, we mapped known CHD4 mutations on our high-resolution structure (Fig. 6, Table 2). Selected sites of mutation are described below. Mutation of residue His1151 to arginine results in a significant reduction of ATPase activity and abolishes chromatin remodelling activity (Kovač et al., 2018). The close proximity of this residue to motif Va (CHD4 residues 1147-1150) makes it likely that the mutation disrupts motif Va function, leading to an uncoupling of the ATPase activity from chromatin remodelling. Similar findings were made for Snf2 where mutation of the tryptophan residue in motif Va resulted in an uncoupling phenotype (Liu et al., 2017). The most frequently mutated residue in endometrial cancer, arginine 1162, is located in the ATPase motif VI. It forms an ‘arginine finger’ that directly interacts with AMP-PNP in our structure. Mutation of Arg1162 to glutamine impairs ATP hydrolysis as suggested by biochemical data (Kovač et al., 2018).

**Table 2.** CHD4 mutations in cancer and Sifrim-Hitz-Weiss syndrome.

|  | Mutated Residue | Location | Predicted effect based on structure | Biochemical observations |
| --- | --- | --- | --- | --- |
| **Cancer** | Cys464Tyr | PHD finger 2 | Disruption of Zn<sup>2+</sup> binding in PHD finger 2 | Reduction in ATPase activity (Kovač et al., 2018) |
|  | Val558Phe | Double chromodomain |  | Reduced ATPase activity (Kovač et al., 2018) |
|  | Arg572Gln | Double chromodomain | Disruption of contact with tracking strand | Reduced DNA binding affinity, Loss of full remodelling activity and ATPase activity (Kovač et al., 2018) |
|  | Leu912Val | ATPase lobe 2 | No prediction made | Reduction of ATPase activity (Kovač et al., 2018) |
|  | His1151Arg | ATPase lobe 2 | In close proximity to motif Va, might disrupt contact of Trp1148 | Reduction of ATPase activity, abolishment of remodelling activity (Kovač et al., 2018) |
|  | Arg1162Gln | ATPase lobe 2, motif VI | Located in ATPase motif VI (arginine finger), Disruption of interaction with ATP | Reduction of ATPase activity (Kovač et al., 2018) |
|  | His1196Tyr | ATPase lobe 2 | Located in the C-terminal bridge region, Removes negative regulation | Speed of chromatin remodelling is increased and better nucleosome centering capability (Kovač et al., 2018) |
|  | Leu1215 | ATPase lobe2/C-terminal bridge | Not located in modelled region |  |
| **Sifrim-Hitz-Weiss syndrome** (Sifrim et al., 2016; Weiss et al., 2016) | Cys467Tyr | PHD finger 2 | Disruption of Zn<sup>2+</sup> binding in PHD finger 2 |  |
|  | Ser851Tyr | ATPase lobe 1 |  |  |
|  | Gly1003Asp | ATPase lobe 2 | Disruption of contact with H3 |  |
|  | Arg1068His | ATPase lobe 2 | Disruption of structural integrity of RecA fold |  |
|  | Arg1127Gln | ATPase lobe 2 | Disruption of contact with DNA minor groove, equivalent arginine residue in SMARCA4 is implicated in “Coffin Siris syndrome” |  |
|  | Trp1148Leu | ATPase lobe 2, motif Va | Disruption of contact with guide strand | Uncoupling of ATPase activity and chromatin remodelling (Liu et al., 2017) |
|  | Arg1173Leu |  | Destabilization |  |
|  | Val1608Ile |  | Not located in modelled region |  |

**Fig. 6.**
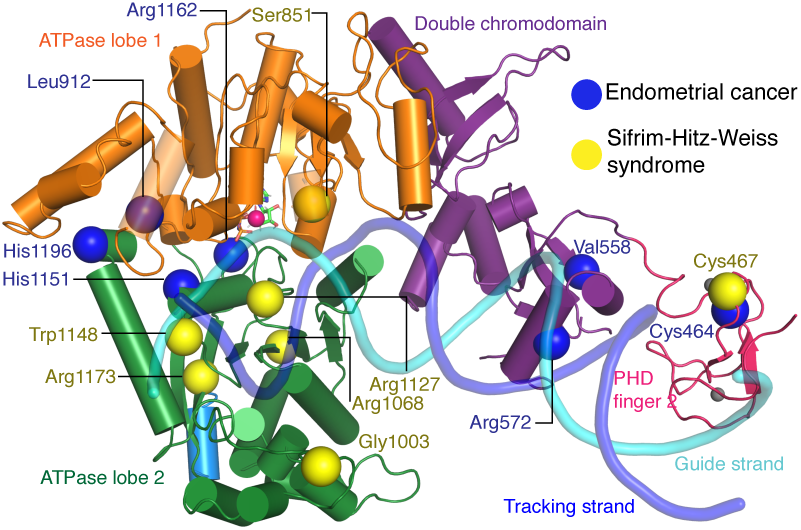
CHD4 mutations in cancer and Sifrim-Hitz-Weiss syndrome. Missense mutations that occur in endometrial cancer (blue spheres) and Sifrim-Hitz-Weiss syndrome (yellow spheres) mapped onto the CHD4 structure. Residue numbering is indicated. Nucleosomal DNA at SHL +2 is shown in a semi-transparent cartoon representation.

### Other disease-related CHD4 mutations

De novo missense mutations in CHD4 are also associated with an intellectual disability syndrome with distinctive dysmorphisms (Sifrim et al., 2016; Weiss et al., 2016). Mutations observed in patients with this syndrome are located in PHD finger 2 (Cys467Tyr) and predominantly in ATPase lobe 2 (Ser851Tyr, Gly1003Asp, Arg1068His, Arg1127Gln, Trp1148Leu, Arg1173Leu, and Val1608Ile). We mapped the sites of these mutations onto our structure (Fig. 6) and predicted the effects of the mutations as far as possible (Table 2).

The Cys467Tyr mutation disrupts coordination of a zinc ion in PHD finger 2. Gly1003 in ATPase lobe2 is located in a loop near H3 alpha helix 1. Deletion of this loop in Chd1 results in a loss of chromatin remodelling activity (Sundaramoorthy et al., 2018). Residue Arg1068 forms a hydrogen bond network with the side chain of Thr1137 and the main chain carbonyl groups of Phe1112 and Gln1119. The Arg1068Cys mutation disrupts this network and is predicted to impair the integrity of the ATPase fold. Mutation of Arg1127 disrupts its interactions with the DNA minor groove (Fig. 2c). The equivalent arginine residue in SMARCA4, which is one of the catalytic subunits of the BAF complex, has been implicated in the rare genetic disorder Coffin-Siris syndrome (Tsurusaki et al., 2012). Trp1148, which is part of ATPase motif Va, contacts the guide strand in a fashion similar to Chd1 and Snf2 (Farnung et al., 2017; Liu et al., 2017) (Fig. 2c). Mutation of this residue uncouples ATP hydrolysis and chromatin remodelling (Liu et al., 2017). Arg1173 inserts into an acidic pocket formed by residues Glu971, Asp1147, and Asp1153. Mutation of the arginine residue to leucine is likely to destabilize ATPase lobe 2 folding.

## Discussion

Here we provide the 3.1 Å resolution cryo-EM structure of human CHD4 engaged with a nucleosome and the 4.0 Å resolution structure of a nucleosome-CHD42 complex that contains two molecules of CHD4. Our structure of the nucleosome-CHD4 complex reveals how a subfamily II CHD remodeller engages with its nucleosomal substrate. We observe a distortion of nucleosomal DNA at SHL +2 in the presence of AMP-PNP. Similar observations were previously made for the Snf2 chromatin remodeller (Li et al., 2019; Liu et al., 2017) in its apo and ADP-bound states.

Our high-resolution structure elucidates the mechanism of chromatin remodelling by capturing an additional enzymatic state. The DNA distortion at SHL +2 that we observed in the AMP-PNP bound state differs from distortions observed previously in the apo and ADP bound state that involved a twist distortion (Li et al., 2019; Winger et al., 2018). This is consistent with a proposed ‘twist defect’ mechanism for chromatin remodelling (Li et al., 2019; Sabantsev et al., 2019). In this model, binding of the ATPase motor at SHL ±2 induces a twist defect in the DNA. Subsequent ATP binding, captured by AMP-PNP and ADP·BeF_3_ structures, then induces closing of the ATPase motor and leads to propagation of the twist defect towards the dyad. It is possible that previous nucleosome-Chd1 structures with ADP·BeF_3_ (Farnung et al., 2017; Sundaramoorthy et al., 2018) also contained the DNA distortion observed here but that their lower resolution prevented its detailed observation. Finally, ATP hydrolysis would reset the remodeller and the enzymatic cycle can resume at the next DNA position.

A major difference between the subfamily II remodeller CHD4 and the subfamily I remodeller Chd1 is that Chd1 induces unwrapping of the terminal nucleosomal DNA, whereas CHD4 does not change the DNA trajectory between SHL -7 to -5. This is likely related to a striking difference in function. Whereas Chd1 functions in euchromatic regions of the genome during active transcription (Skene et al., 2014), CHD4 plays a central role in the establishment and maintenance of repressive genome regions. Consistent with these findings, DNA unwrapping should be prevented in stable heterochromatic regions. It is likely that the evolution of auxiliary domains in different CHD subfamilies led to these different functionalities. In particular, the DNA-binding region in Chd1 or the PHD fingers in CHD4 alter the functional properties of these chromatin remodellers, with the former working on active genes, and the latter often functioning in gene repression. Our structure also helps to define how causative disease mutations impair CHD4 function. Mutations in disease phenotypes are able to disrupt DNA binding, impede ATP hydrolysis, or uncouple ATP hydrolysis and DNA translocation. The structure rationalizes the effects of CHD4 mutations in cancer and intellectual disability syndromes on chromatin remodelling. It also helps in understanding disease phenotypes of other chromatin remodellers such as the BAF complex that shows a related domain architecture for its ATPase motor. Due to its high resolution, the structure may also guide drug discovery using chromatin remodellers as targets.

## ACKNOWLEDGEMENTS

We thank past and present members of the Cramer laboratory. We thank C. Oberthür for help with protein purification, U. Neef for insect cell maintenance, A. Sawicka for providing cDNA, and S.M. Vos for valuable input and critical reading of the manuscript. P.C. was supported by the Deutsche Forschungsgemeinschaft (SFB1064, SFB860), the European Research Council Advanced Investigator Grant TRANSREGULON (grant agreement No. 693023), and the Volkswagen Foundation.

## AUTHOR CONTRIBUTIONS

L.F. designed and carried out all experiments. L.F. and M.O. performed model building. L.F. designed research. P.C. supervised research. L.F. and M.O. generated figures. L.F. and P.C. prepared the manuscript, with input from M.O.

## DATA AVAILABILITY

The cryo-EM reconstructions and final models were deposited with the Electron Microscopy Data Bank (accession codes EMD-10058 and EMD-10059) and with the Protein Data Bank (accession code 6RYR and 6RYU).

## Methods

### Preparation of CHD4

*H. sapiens* CHD4 (Uniprot Accession code Q14839-1) was amplified from human cDNA using the following ligation-independent cloning (LIC) compatible primer pair (Forward primer: 5’-TAC TTC CAA TCC AAT GCA ATG GCG TCG GGC CTG-3’, reverse primer: 5’-TTA TCC ACT TCC AAT GTT ATT ACT GCT GCT GGG CTA CCT G-3’). The PCR product containing CHD4 was cloned into a modified pFastBac vector (a gift from S. Gradia, UC Berkeley, vector 438-C, Addgene: 55220) via LIC. The CHD4 construct contains a N-terminal 6xHis tag, followed by a MBP tag, a 10x Asn linker sequence, and a tobacco etch virus protease cleavage site. All sequences were verified by Sanger sequencing.

The CHD4 plasmid (500 ng) was electroporated into DH10EMBacY cells (Geneva Biotech) to generate a bacmid encoding full-length *H. sapiens* CHD4. Bacmids were subsequently selected and prepared from positive clones using blue/white selection and isopropanol precipitation. V0 and V1 virus production was performed as previously described49. Hi5 cells (600 ml) grown in ESF-921 media (Expression Systems) were infected with 200 µl of V1 virus for protein expression. The cells were grown for 72 h at 27 °C. Cells were harvested by centrifugation (238g, 4 °C, 30 min) and resuspended in lysis buffer (300 mM NaCl, 20 mM Na·HEPES pH 7.4, 10% (v/v) glycerol, 1 mM DTT, 30 mM imidazole pH 8.0, 0.284 µg mL ^-1^ leupeptin, 1.37 µg mL^-1^ pepstatin A, 0.17 mg mL^-1^ PMSF, 0.33 mg mL^-1^ benzamidine). The cell resuspension was frozen and stored at -s80 °C.

*H. sapiens* CHD4 was purified at 4 °C. Frozen cell pellets were thawed and lysed by sonication. Lysates were cleared by two centrifugation steps (18,000g, 4 °C, 30 min and 235,000g, 4 °C, 60 min). The supernatant containing CHD4 was filtered using 0.8-µm syringe filters (Millipore). The filtered sample was applied onto a GE HisTrap HP 5 ml (GE Healthcare), pre-equilibrated in lysis buffer. After sample application, the column was washed with 10 CV lysis buffer, 5 CV high salt buffer (1 M NaCl, 20 mM Na·HEPES pH 7.4, 10% (v/v) glycerol, 1 mM DTT, 30 mM imidazole pH 8.0, 0.284 µg mL^-1^ leupeptin, 1.37 µg mL^-1^ pepstatin A, 0.17 mg mL^-1^ PMSF, 0.33 mg mL^-1^ benzamidine), and 5 CV lysis buffer. The protein was eluted with a gradient of 0–100% elution buffer (300 mM NaCl, 20 mM Na·HEPES p 7.4, 10% (v/v) glycerol, 1 mM DTT, 500 mM imidazole pH 8.0, 0.284 µg mL^-1^ leupeptin, 1.37 µg mL^-1^ pepstatin A, 0.17 mg ml^-1^ PMSF, 0.33 mg mL^-1^ benzamidine). Peak fractions were pooled and dialysed for 16 h against 600 ml dialysis buffer (300 mM NaCl, 20 mM Na·HEPES pH 7.4, 10% (v/v) glycerol, 1 mM DTT, 30 mM imidazole) in the presence of 2 mg His6-TEV protease. The dialysed sample was applied to a GE HisTrap HP 5 ml. The flow-through containing CHD4 was concentrated using an Amicon Millipore 15 ml 50,000 MWCO centrifugal concentrator. The concentrated CHD4 sample was applied to a GE S200 16/600 pg size exclusion column, pre-equilibrated in gel filtration buffer (300 mM NaCl, 20 mM Na·HEPES pH 7.4, 10% (v/v) glycerol, 1 mM DTT). Peak fractions were concentrated to 40 µM, aliquoted, flash frozen, and stored at -80 °C. Typical yields of *H. sapiens* CHD4 from 1.2 L of Hi5 insect cell culture are 2-4 mg.

### Nucleosome Preparation

*Xenopus laevis* histones were expressed and purified as described (Dyer et al., 2003; Farnung et al., 2017). DNA fragments for nucleosome reconstitution were generated by PCR essentially as described (Farnung et al., 2018). A vector containing the Widom 601 sequence was used as a template for PCR. Super-helical locations are assigned based on previous publications (Farnung et al., 2018, 2017; Kujirai et al., 2018; Sundaramoorthy et al., 2018), assuming potential direction of transcription from negative to positive SHLs. Large-scale PCR reactions were performed with two PCR primers (forward primer: TGT TGG ATG TTT TAT AAT TGA GTG GGT TCC TGT TAT TCC TAG TAA TCA ATC AGT GCC TAT CGA TGT ATA TAT CTG ACA CGT GCC T, reverse primer: CCC CAT CAG AAT CCC GGT GCC G) at a scale of 25 mL. Nucleosome core particle reconstitution was performed using the salt-gradient dialysis method (Dyer et al., 2003). Quantification of the reconstituted nucleosome was achieved by measuring absorbance at 280 nm. Molar extinction coefficients were determined for protein and nucleic acid components and were summed to yield a molar extinction coefficient for the reconstituted extended nucleosome.

### Reconstitution of nucleosome-CHD4 complex

Reconstituted nucleosome core particles and CHD4 were mixed at a molar ratio of 1:2. AMP-PNP was added at a final concentration of 1 mM and the sample was incubated for 10 minutes on ice. After 10 minutes compensation buffer was added to a final buffer concentration of 30 mM NaCl, 3 mM MgCl_2_, 20 mM Na·HEPES pH 7.5, 4% (v/v) glycerol, 1 mM DTT. The sample was applied to a Superose 6 Increase 3.2/300 column equilibrated in gel filtration buffer (30 mM NaCl, 3 mM MgCl_2_, 20 mM Na·HEPES pH 7.5, 5% (v/v) glycerol, 1 mM DTT). The elution was fractionated in 50 µL fractions and peak fractions were analysed by SDS-PAGE. Relevant fractions containing nucleosome core particle and CHD4 were selected and cross-linked with 0.1% (v/v) glutaraldehyde. The crosslinking reaction was performed for 10 min on ice and subsequently quenched for 10 min using a final concentration of 2 mM lysine and 8 mM aspartate. The sample was transferred to a Slide-A-Lyzer MINI Dialysis Unit 20,000 MWCO (Thermo Scientific), and dialysed for 4 h against 600 ml dialysis buffer (30 mM NaCl, 3 mM MgCl_2_, 20 mM Na·HEPES pH 7.4, 20 mM Tris·HCl pH 7.5, 1 mM DTT). The sample was subsequently concentrated using a Vivaspin 500 ultrafiltration centrifugal concentrator (Sartorius) to a final concentration of 200-300 µM.

### Cryo-EM analysis and image processing

The nucleosome-CHD4 sample was applied to R2/2 gold grids (Quantifoil). The grids were glow-discharged for 100 s before sample application of 2 µl on each side of the grid. The sample was subsequently blotted for 8.5 s (Blot force 5) and vitrified by plunging into liquid ethane with a Vitrobot Mark IV (FEI Company) operated at 4 °C and 10% humidity. Cryo-EM data were acquired on a Titan Krios transmission electron microscope (FEI/Thermo) operated at 300 keV, equipped with a K2 summit direct detector (Gatan) and a GIF Quantum energy filter. Automated data acquisition was carried out using FEI EPU software at a nominal magnification of 130,000x in nanoprobe EF-TEM mode. Image stacks of 40 frames were collected in counting mode over 10 s. The dose rate was 4.3-4.5 e^-^ per Å^2^ per s for a total dose of 43-45 e^-^ per Å^2^. A total of 3,904 image stacks were collected.

Micrograph frames were stacked and processed. All micrographs were CTF and motion corrected using Warp (Tegunov and Cramer, 2018). Particles were picked using an in-house trained instance of the neural network BoxNet2 of Warp, yielding 650,598 particle positions. Particles were extracted with a box size of 300^2^ pixel and normalized. Image processing was performed with RELION 3.0-beta 2 (Zivanov et al., 2018). Using a 30 Å low-pass filtered ab initio model generated in cryoSPARC from 1,679 particles (Supplementary Fig. 2c) we performed one round of 3D classification of all particle images with image alignment. One class with defined density for the nucleosome-CHD4 complex was selected for a second round of classification. The second round of classification resulted in two classes with one copy of CHD4 bound to the nucleosome. The respective classes were selected and 3D refined. The refined nucleosome-CHD4 model was subsequently CTF refined and the beam tilt was estimated based on grouping of beam tilt classes according to their exposure positions. The CTF refined particles were submitted to one additional round of masked 3D classification without image alignment. The mask encompassed CHD4. The most occupied class from this classification was subsequently CTF-refined. The final particle reconstruction was obtained from a 3D refinement with a mask that encompasses the entire nucleosome-CHD4 complex.

The nucleosome-CHD4 reconstruction was obtained from 89,623 particles with an overall resolution of 3.1 Å (gold-standard Fourier shell correlation 0.143 criterion). The final map was sharpened with a B-factor of -36 Å^2^. Additionally, the second round of 3D classification yielded a class with a nucleosome-CHD4_2_ complexes. The particles were subsequently classified and refined. The resulting reconstruction with 40,233 particles had an overall resolution of 4.0 Å (gold-standard Fourier shell correlation 0.143 criterion). The final map was sharpened with a B-factor of -86 Å^2^. Local resolution estimates for both structures were determined using the built-in RELION tool.

### Model building

Crystal structures of the X. laevis nucleosome with the Widom 601 sequence (Vasudevan et al., 2010) (PDB code 3LZ0) and the double chromodomain of CHD4 (PDB code 4O9I) were placed into the density of the nucleosome-CHD4 complex as rigid bodies using UCSF Chimera. The protein sequence of the ATPase motor of CHD4 (residues 706-1196) was ‘one-to-one threaded’ using the ATPase motor of *S. cerevisiae* Chd1 (PDB code 5O9G) as a template by employing Phyre2 (Kelley et al., 2015). The threaded model was placed into the density as a rigid body using UCSF Chimera (Goddard et al., 2018). Additional density belonging to helical extensions and loops present in the ATPase motor region were modelled de novo.

The nucleosome structure, double chromodomain structure, and ATPase motor model were adjusted manually in COOT (version 0.9-pre) (Emsley et al., 2010). The structure of PHD finger 2 (Mansfield et al., 2011) was then manually placed into the remaining, weaker density next to the double chromodomain and rigid-body docked (Supplementary Fig. 3). Additional structural elements such as the H4 tail, the C-terminal bridge and loop regions of CHD4 were built using COOT. AMP-PNP and a coordinated Mg2+ ion were placed into the corresponding density. AMP-PNP was derived from the monomer library in COOT. The high resolution of our reconstruction enabled us to model several DNA-interacting side chains in two alternative conformations. The complete model was real-space refined in PHENIX (Afonine et al., 2018) with global minimization, local rotamer fitting, morphing and simulated annealing. To model the nucleosome-CHD4_2_ complex, the CHD4 model was duplicated and the second copy was rigid body docked into the additional density using UCSF ChimeraX (Goddard et al., 2018). The resulting structure was real space refined in PHENIX with global minimization, local rotamer fitting, morphing and simulated annealing.

### Figure generation

Figures were generated using PyMol (version 2.2.2) and UCSF ChimeraX.

### Latex Template

Ricardo Henriques biorxiv template was used to render this pre-print (https://www.overleaf.com/latex/templates/henriqueslab-biorxiv-template/nyprsybwffws).

## Supplementary Figures

**Supplementary Figure 1.**
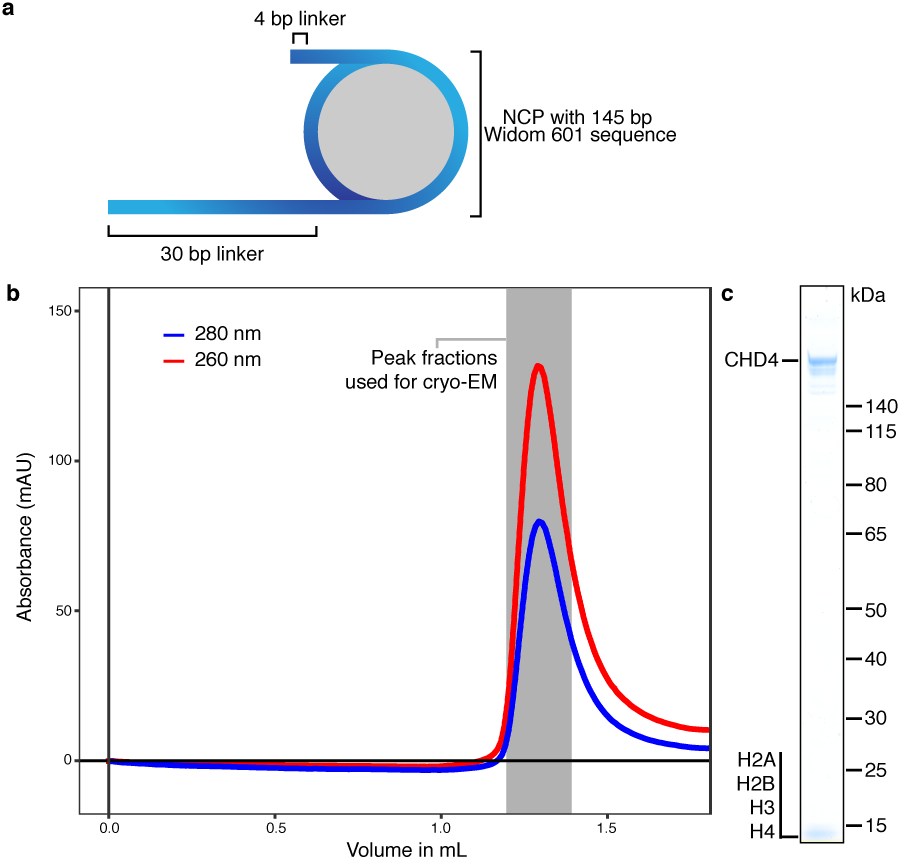
Formation of the nucleosome-CHD4 complex. a, Schematic of DNA construct to form nucleosome-CHD4 complex. Extranucleosomal DNA length is indicated. b, Formation of the nucleosome-CHD4 complex on a Superose 6 Increase 3.2/30 size exclusion chromatography column. Red and blue curve shows absorption at 260 nm and 280 nm milli absorption units, respectively. c, SDS-PAGE gel with peak fraction containing the formed nucleosome-CHD4 complex.

**Supplementary Figure 2.**
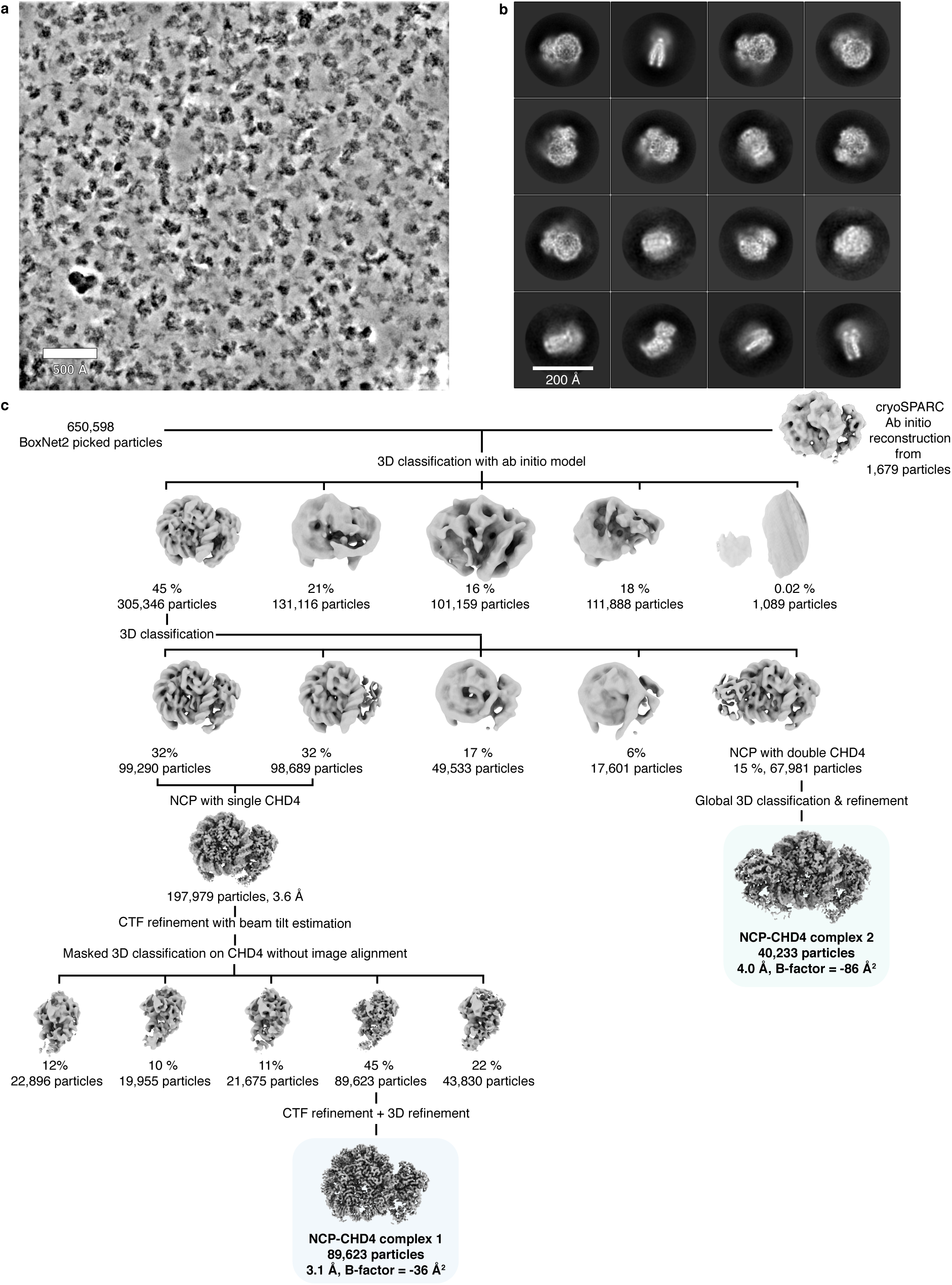
Cryo-EM structure determination. a, Representative micrograph of data collection. The micrograph was denoised using Warp(Tegunov and Cramer, 2018). Scale bar with a length of 500 Å is shown. b, 2D classes of single copy CHD4 bound to a nucleosome. Scale bar with a length of 200 Å is shown. c, Classification tree employed to obtain cryo-EM density of CHD4 bound to a nucleosome. Particle numbers and class distribution percentages are indicated. Final reconstructions are highlighted.

**Supplementary Figure 3.**
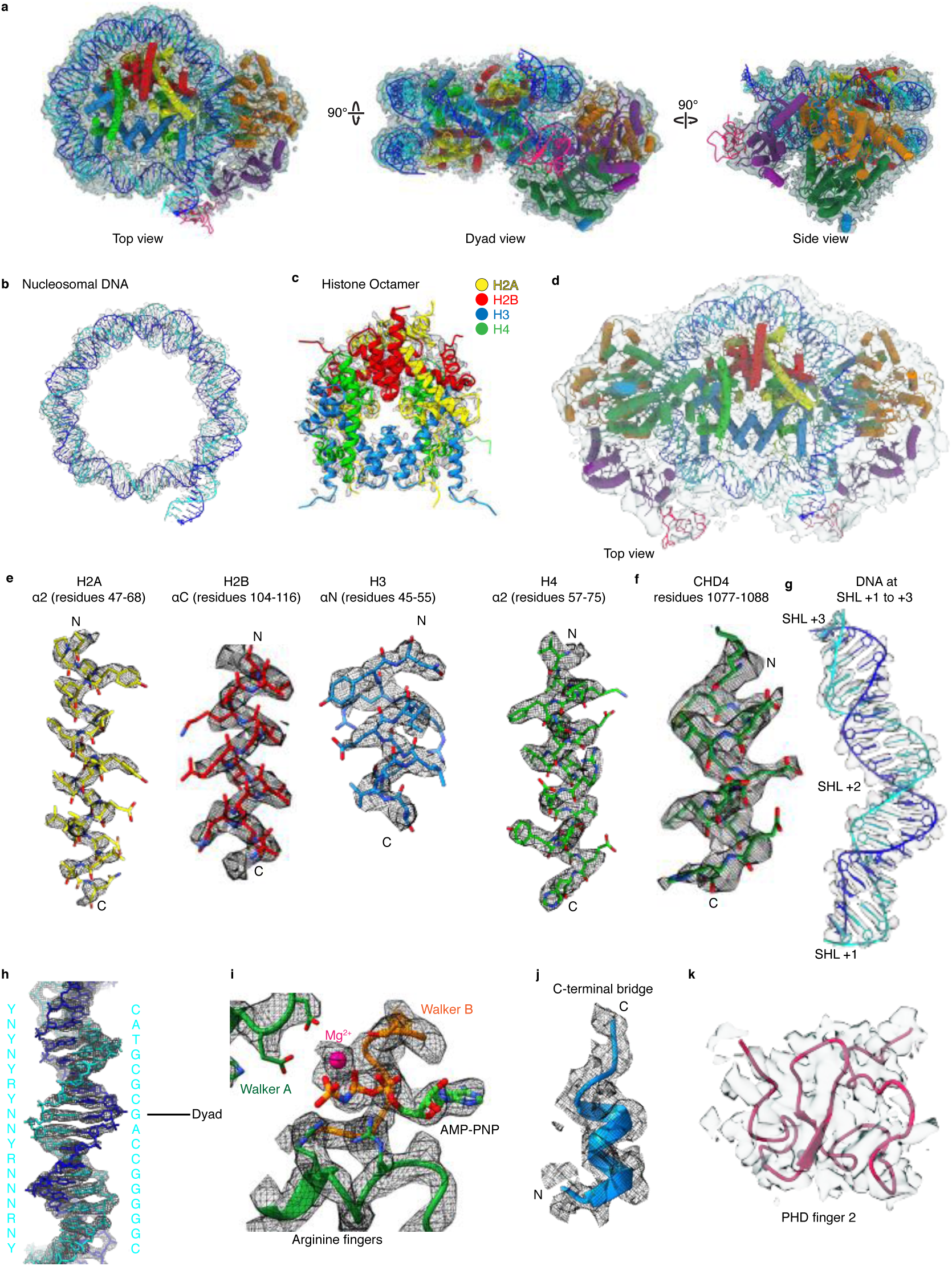
Cryo-EM densities. a, Cartoon model of CHD4-nucleosome structure with corresponding post-processed Coulomb potential map shown in silver. b, Nucleosomal DNA with Coulomb potential map. c, Histone octamer with Coulomb potential map d, Cartoon model of two copies of CHD4 engaged with the nucleosome and corresponding Coulomb potential map. e, Representative density of histone residues. f, Representative density of CHD4 residues. g, Coulomb potential map of density near DNA at SHL +2. h, DNA density around dyad axis with fitted DNA model. Base identities used to fit register and directionality are indicated on the left. N, R, and Y indicate any nucleotide, purine, or pyrimidine, respectively. Matching sequence provided on the right. i, Active site density with fitted AMP-PNP and coordinated Mg2+ ion. j, Density of C-terminal bridge helix. k, Cartoon model of PHD finger 2 with corresponding local resolution filtered Coulomb potential map.

**Supplementary Figure 4.**
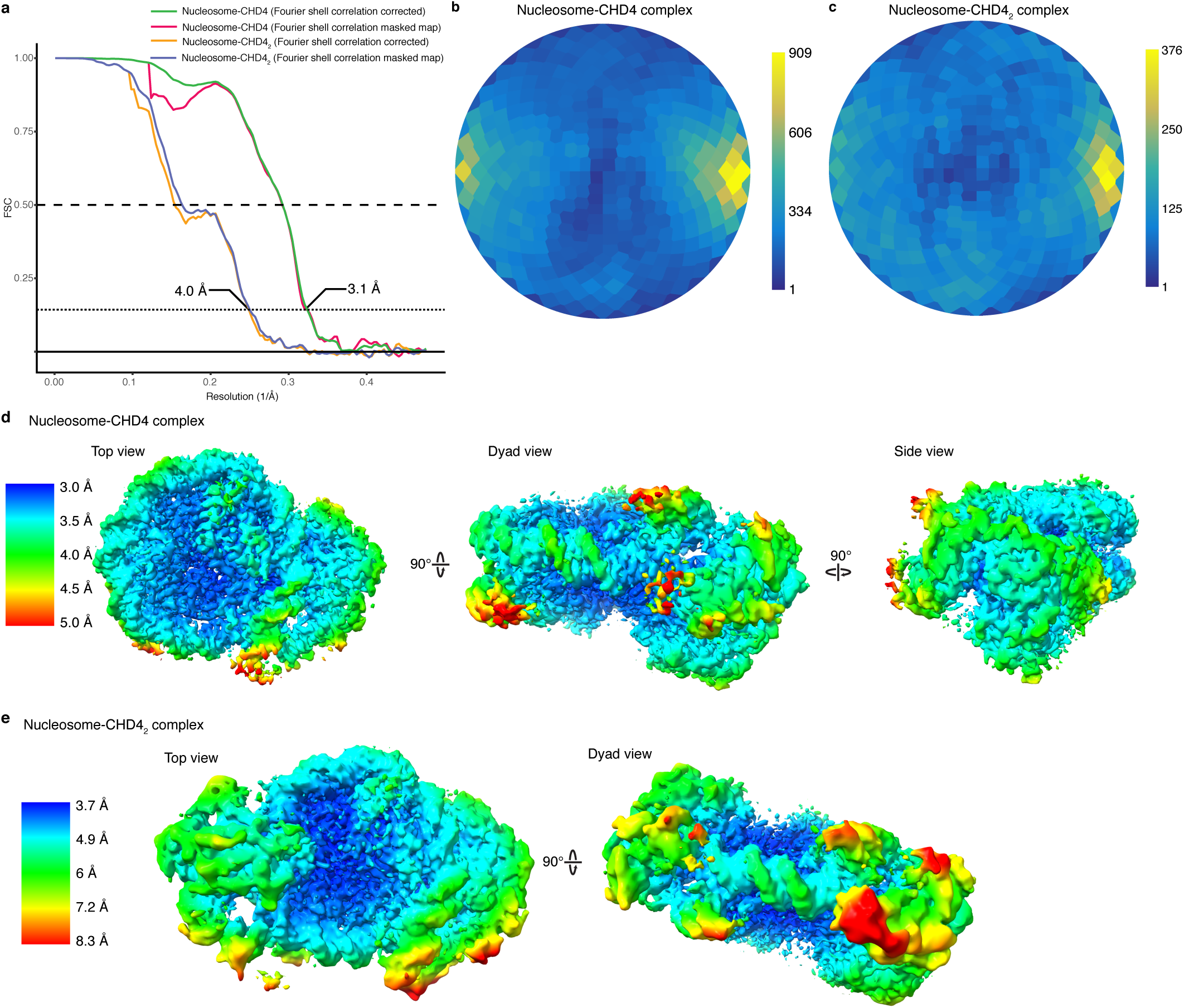
Data quality and metrics. a, FSC curves. b, Angular distribution plots. c, Local resolution of CHD4 structures. Densities are coloured according to resolution as indicated.

**Supplementary Figure 5.**
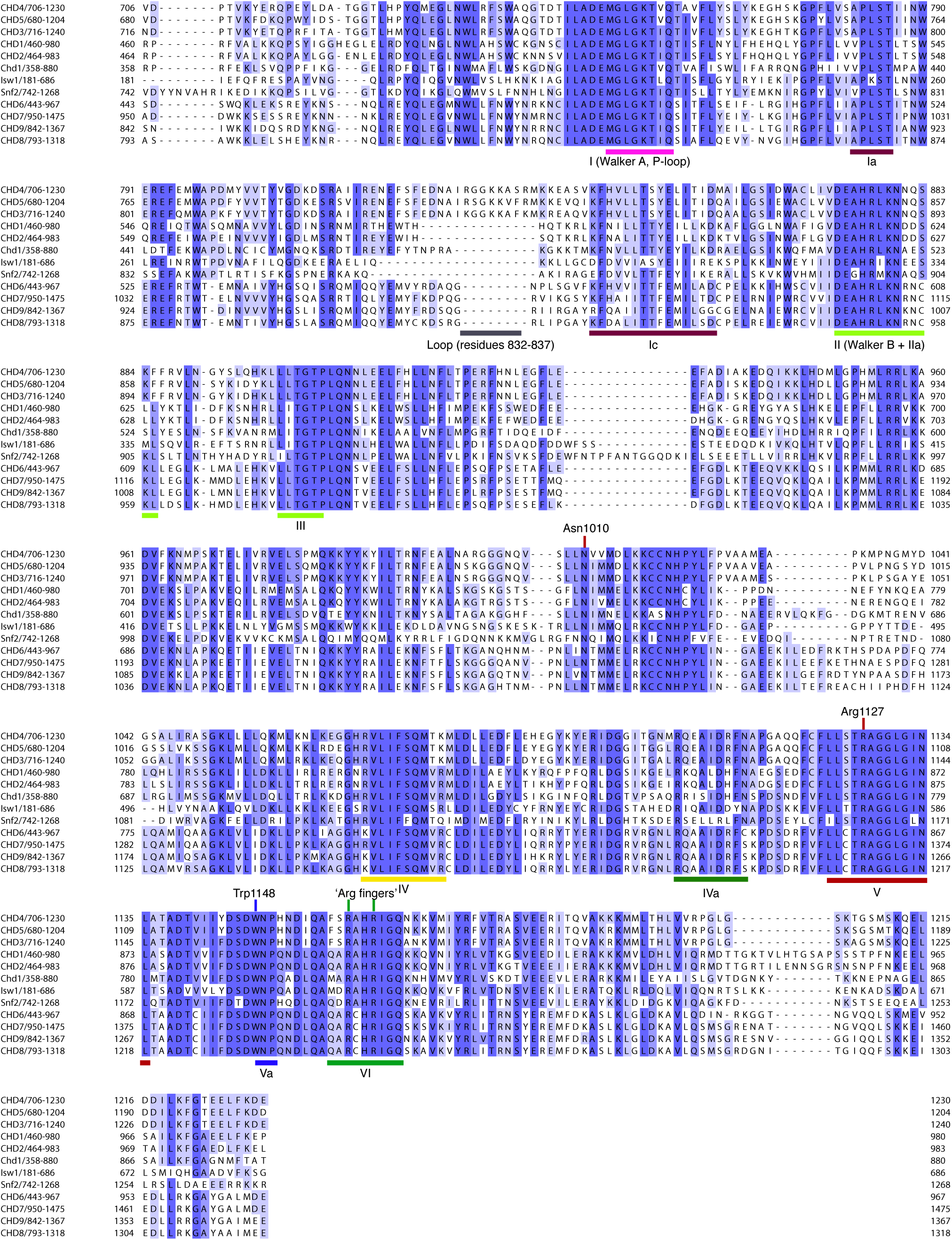
Comparison of CHD4 with Chd1 and other chromatin remodellers. Sequence alignment of ATPase regions in *H. sapiens* CHD4 (706-1230), CHD5 (680-1204), CHD3 (716-1240), CHD1 (460-980), CHD2 (464-983), *S. cerevisiae* Chd1 (358-880), *S. cerevisiae* Isw1 (181-686), *S. cerevisiae* Snf2 (742-1268), H. sapiens CHD6 (443-967), CHD7 (950-1475), CHD9 (842-1367), and CHD8 (793-1318). Important elements and ATPase motifs are indicated. Sequence coloured according to identity. Dark and light shades of blue indicate high and low conservation, respectively. Alignment generated with MAFFT (Katoh and Standley, 2013) and visualized using JalView (Waterhouse et al., 2009).

## References

Afonine PV, Poon BK, Read RJ, Sobolev OV, Terwilliger TC, Urzhumtsev A, Adams PD. 2018. Real-space refinement in PHENIX for cryo-EM and crystallography. Acta Crystallogr Sect D Struct Biology 74:531–544. doi:10.1107/s2059798318006551

Armache J-P, Gamarra N, Johnson SL, Leonard JD, Wu S, Narlikar GJ, Cheng Y. 2019. Electron cryo-microscopy structures of remodeler-nucleosome intermediates suggest allosteric control through the nucleosome. Biorxiv 550970. doi:10.1101/550970

Bornelöv S, Reynolds N, Xenophontos M, Gharbi S, Johnstone E, Floyd R, Ralser M, Signolet J, Loos R, Dietmann S, Bertone P, Hendrich B. 2018. The Nucleosome Remodeling and Deacetylation Complex Modulates Chromatin Structure at Sites of Active Transcription to Fine-Tune Gene Expression. Mol Cell 71:56-72.e4. doi:10.1016/j.molcel.2018.06.003

Brandani GB, Niina T, Tan C, Takada S. 2018. DNA sliding in nucleosomes via twist defect propagation revealed by molecular simulations. Nucleic Acids Res 46:gky158.. doi:10.1093/nar/gky158

Brandani GB, Takada S. 2018. Chromatin remodelers couple inchworm motion with twist-defect formation to slide nucleosomal DNA. Plos Comput Biol 14:e1006512. doi:10.1371/journal.pcbi.1006512

Burgold T, Barber M, Kloet S, Cramard J, Gharbi S, Floyd R, Kinoshita M, Ralser M, Vermeulen M, Reynolds N, Dietmann S, Hendrich B. 2019. The Nucleosome Remodelling and Deacetylation complex suppresses transcriptional noise during lineage commitment. Embo J. doi:10.15252/embj.2018100788

Clapier CR, Iwasa J, Cairns BR, Peterson CL. 2017. Mechanisms of action and regulation of ATP-dependent chromatin-remodelling complexes. Nat Rev Mol Cell Bio 18:407–422. doi:10.1038/nrm.2017.26

Dyer PN, Edayathumangalam RS, White CL, Bao Y, nivas Chakravarthy, Muthurajan UM, Luger K. 2003. Methods in Enzymology 375:23–44. doi:10.1016/s0076-6879(03)75002-2

Emsley P, Lohkamp B, Scott W, Cowtan K. 2010. Features and development of Coot. Acta Crystallogr Sect D Biological Crystallogr 66:486–501. doi:10.1107/s0907444910007493

Farnung L, Vos SM, Cramer P. 2018. Structure of transcribing RNA polymerase II-nucleosome complex. Nat Commun 9:5432. doi:10.1038/s41467-018-07870-y

Farnung L, Vos SM, Wigge C, Cramer P. 2017. Nucleosome–Chd1 structure and implications for chromatin remodelling. Nature 550:539. doi:10.1038/nature24046

Gatchalian J, Wang X, Ikebe J, Cox KL, Tencer AH, Zhang Y, Burge NL, Di L, Gibson MD, Musselman CA, Poirier MG, Kono H, Hayes JJ, Kutateladze TG. 2017. Accessibility of the histone H3 tail in the nucleosome for binding of paired readers. Nat Commun 8:1489. doi:10.1038/s41467-017-01598-x

Getz G, Gabriel SB, Cibulskis K, Lander E, Sivachenko A, Sougnez C, Lawrence M, Kandoth C, Dooling D, Fulton R, Fulton L, Kalicki-Veizer J, McLellan MD, O’Laughlin M, Schmidt H, Wilson RK, Ye K, Ding L, Mardis ER, Ally A, Balasundaram M, Birol I, Butterfield YS, Carlsen R, Carter C, Chu A, Chuah E, Chun H-JE, Dhalla N, Guin R, Hirst C, Holt RA, Jones SJ, Lee D, Li HI, Marra MA, Mayo M, Moore RA, Mungall AJ, Plettner P, Schein JE, Sipahimalani P, Tam A, Varhol RJ, Robertson GA, Cherniack AD, Pashtan I, Saksena G, Onofrio RC, Schumacher SE, Tabak B, Carter SL, Hernandez B, Gentry J, Salvesen HB, Ardlie K, Winckler W, Beroukhim R, Meyerson M, Hadjipanayis A, Lee S, Mahadeshwar HS, Park P, Protopopov A, Ren X, Seth S, Song X, Tang J, Xi R, Yang Lixing, Zeng D, Kucherlapati R, Chin L, Zhang J, Auman TJ, Balu S, Bodenheimer T, Buda E, Hayes ND, Hoyle AP, Jefferys SR, Jones CD, Meng S, Mieczkowski PA, Mose LE, Parker JS, Perou CM, Roach J, Shi Y, Simons JV, Soloway MG, Tan D, Topal MD, Waring S, Wu J, Hoadley KA, Baylin SB, Bootwalla MS, Lai PH, Jr TJ, Berg DJ, Weisenberger DJ, Laird PW, Shen H, Cho J, DiCara D, Frazer S, Heiman D, Jing R, Lin P, Mallard W, Stojanov P, Voet D, Zhang H, Zou L, Noble M, Reynolds SM, Shmulevich I, man Aksoy, Antipin Y, Ciriello G, Dresdner G, Gao J, Gross B, Jacobsen A, Ladanyi M, Reva B, Sander C, Sinha R, Sumer OS, Taylor BS, Cerami E, Weinhold N, Schultz N, Shen R, Benz S, Goldstein T, Haussler D, Ng S, Szeto C, Stuart J, Benz CC, Yau C, Zhang W, Annala M, Broom BM, Casasent TD, Ju Z, Liang H, Liu G, Lu Y, Unruh AK, Wakefield C, Weinstein JN, Zhang N, Liu Y, Broaddus R, Akbani R, Mills GB, Adams C, Barr T, Black AD, Bowen J, Deardurff J, Frick J, Gastier-Foster JM, Grossman T, Harper HA, Hart-Kothari M, Helsel C, Hobensack A, Kuck H, Kneile K, Leraas KM, Lichtenberg TM, McAllister C, Pyatt RE, Ramirez NC, Tabler TR, Vanhoose N, White P, Wise L, Zmuda E, Barnabas N, Berry-Green C, Blanc V, Boice L, Button M, Farkas A, Green A, MacKenzie J, Nicholson D, Kalloger SE, Gilks BC, Karlan BY, Lester J, Orsulic S, Borowsky M, Cadungog M, Czerwinski C, Huelsenbeck-Dill L, Iacocca M, Petrelli N, Rabeno B, Witkin G, Nemirovich-Danchenko E, Potapova O, Rotin D, Berchuck A, Birrer M, DiSaia P, Monovich L, Curley E, Gardner J, Mallery D, Penny R, Dowdy SC, Winterhoff B, Dao L, Gostout B, Meuter A, Teoman A, Dao F, Olvera N, Bogomolniy F, Garg K, Soslow RA, Levine DA, Abramov M, Bartlett J, Kodeeswaran S, Parfitt J, Moiseenko F, Clarke BA, Goodman MT, Carney ME, Matsuno RK, Fisher J, Huang M, Rathmell KW, Thorne L, Le L, Dhir R, Edwards R, Elishaev E, Zorn K, Goodfellow PJ, Mutch D, Kahn AB, Bell DW, Pollock PM, Wang C, Wheeler DA, Shinbrot E, Ayala B, Chu AL, Jensen MA, Kothiyal P, Pihl TD, Pontius J, Pot DA, Snyder EE, nivasan D, Shaw KR, Sheth M, Davidsen T, Ferguson GL, Demchok JA, Yang Liming, Guyer MS, Ozenberger BA, Sofia HJ. 2013. Integrated genomic characterization of endometrial carcinoma. Nature 497:67. doi:10.1038/nature12113

Gnanapragasam M, Scarsdale NJ, Amaya ML, Webb HD, Desai MA, Walavalkar NM, Wang S, Zhu S, Ginder GD, Williams DC. 2011. p66a–MBD2 coiled-coil interaction and recruitment of Mi-2 are critical for globin gene silencing by the MBD2–NuRD complex. Proc National Acad Sci 108:7487–7492. doi:10.1073/pnas.1015341108

Goddard TD, Huang CC, Meng EC, Pettersen EF, Couch GS, Morris JH, Ferrin TE. 2018. UCSF ChimeraX: Meeting modern challenges in visualization and analysis. Protein Sci 27:14–25. doi:10.1002/pro.3235

Hauk G, McKnight JN, Nodelman IM, Bowman GD. 2010. The Chromodomains of the Chd1 Chromatin Remodeler Regulate DNA Access to the ATPase Motor. Mol Cell 39:711–23. doi:10.1016/j.molcel.2010.08.012

Katoh K, Standley DM. 2013. MAFFT Multiple Sequence Alignment Software Version 7: Improvements in Performance and Usability. Mol Biol Evol 30:772–780. doi:10.1093/molbev/mst010

Kehle J, Beuchle D, Treuheit S, Christen B, Kennison JA, Bienz M, Müller J. 1998. dMi-2, a Hunchback-Interacting Protein That Functions in Polycomb Repression. Science 282:1897–1900. doi:10.1126/science.282.5395.1897

Kelley LA, Mezulis S, Yates CM, Wass MN, Sternberg MJ. 2015. The Phyre2 web portal for protein modeling, prediction and analysis. Nat Protoc 10:845–858. doi:10.1038/nprot.2015.053

Kovač K, Sauer A, Mačinković I, Awe S, Finkernagel F, Hoffmeister H, Fuchs A, Müller R, Rathke C, Längst G, Brehm A. 2018. Tumour-associated missense mutations in the dMi-2 ATPase alters nucleosome remodelling properties in a mutation-specific manner. Nat Commun 9:2112. doi:10.1038/s41467-018-04503-2

Kujirai T, Ehara H, Fujino Y, Shirouzu M, Sekine S, Kurumizaka H. 2018. Structural basis of the nucleosome transition during RNA polymerase II passage. Science 362:eaau9904. doi:10.1126/science.aau9904

Kuzmichev A, Nishioka K, Erdjument-Bromage H, Tempst P, Reinberg D. 2002. Histone methyltransferase activity associated with a human multiprotein complex containing the Enhancer of Zeste protein. Gene Dev 16:2893–2905. doi:10.1101/gad.1035902

Kwan A, Gell DA, Verger A, Crossley M, Matthews JM, Mackay JP. 2003. Engineering a Protein Scaffold from a PHD Finger. Structure 11:803–813. doi:10.1016/s0969-2126(03)00122-9

Längst G, Manelyte L. 2015. Chromatin Remodelers: From Function to Dysfunction. Genes-basel 6:299–324. doi:10.3390/genes6020299

Larsen D, Poinsignon C, Gudjonsson T, Dinant C, Payne MR, Hari FJ, Danielsen JM, Menard P, Sand J, Stucki M, Lukas C, Bartek J, Andersen JS, Lukas J. 2010. The chromatin-remodeling factor CHD4 coordinates signaling and repair after DNA damage. J Cell Biology 190:731–740. doi:10.1083/jcb.200912135

Li Meijing, Xia X, Tian Y, Jia Q, Liu X, Lu Y, Li Ming, Li X,Chen Z. 2019. Mechanism of DNA translocation underlying chromatin remodelling by Snf2. Nature 567:409–413. doi:10.1038/s41586-019-1029-2

Liang Z, Brown KE, Carroll T, Taylor B, Vidal I, Hendrich B, Rueda D, Fisher AG, Merkenschlager M. 2017. A high-resolution map of transcriptional repression. Elife 6:e22767. doi:10.7554/elife.22767

Liu X, Li M, Xia X, Li X, Chen Z. 2017. Mechanism of chromatin remodelling revealed by the Snf2-nucleosome structure. Nature 544:440. doi:10.1038/nature22036

Lowary P., Widom J. 1998. New DNA sequence rules for high affinity binding to histone octamer and sequence-directed nucleosome positioning11Edited by T. Richmond. J Mol Biol 276:19–42. doi:10.1006/jmbi.1997.1494

Mansfield RE, Musselman CA, Kwan AH, Oliver SS, Garske AL, Davrazou F, Denu JM, Kutateladze TG, Mackay JP. 2011. Plant Homeodomain (PHD) Fingers of CHD4 Are Histone H3-binding Modules with Preference for Unmodified H3K4 and Methylated H3K9. J Biol Chem 286:11779–11791. doi:10.1074/jbc.m110.208207

Nodelman IM, Bleichert F, Patel A, Ren R, Horvath KC, Berger JM, Bowman GD. 2017. Interdomain Communication of the Chd1 Chromatin Remodeler across the DNA Gyres of the Nucleosome. Mol Cell 65:447-459.e6. doi:10.1016/j.molcel.2016.12.011

Ostapcuk V, Mohn F, Carl SH, Basters A, Hess D, Iesmantavicius V, Lampersberger L, Flemr M, Pandey A, Thomä NH, Betschinger J, Bühler M. 2018. Activity-dependent neuroprotective protein recruits HP1 and CHD4 to control lineage-specifying genes. Nature 557:739–743. doi:10.1038/s41586-018-0153-8

Polo SE, Kaidi A, Baskcomb L, Galanty Y, Jackson SP. 2010. Regulation of DNA-damage responses and cell-cycle progression by the chromatin remodelling factor CHD4. Embo J 29:3130–3139. doi:10.1038/emboj.2010.188

Sabantsev A, Levendosky RF, Zhuang X, Bowman GD, Deindl S. 2019. Direct observation of coordinated DNA movements on the nucleosome during chromatin remodelling. Nat Commun 10:1720. doi:10.1038/s41467-019-09657-1

Schalch T, Duda S, Sargent DF, Richmond TJ. 2005. X-ray structure of a tetranucleosome and its implications for the chromatin fibre. Nature 436:138. doi:10.1038/nature03686

Schindler U, Beckmann H, Cashmore AR. 1993. HAT3.1, a novel Arabidopsis homeodomain protein containing a conserved cysteine-rich region. Plant J 4:137–150. doi:10.1046/j.1365-313x.1993.04010137.x

Sifrim A, Hitz M-P, Wilsdon A, Breckpot J, Turki SH, Thienpont B, McRae J, Fitzgerald TW, Singh T, Swaminathan G, Prigmore E, Rajan D, Abdul-Khaliq H, Banka S, Bauer UM, Bentham J, Berger F, Bhattacharya S, Bu’Lock F, Canham N, Colgiu I-G, Cosgrove C, Cox H, Daehnert I, Daly A, Danesh J, Fryer A, Gewillig M, Hobson E, Hoff K, Homfray T, Study I, Kahlert A-K, Ketley A, Kramer H-H, Lachlan K, Lampe A, Louw JJ, Manickara A, Manase D, McCarthy KP, Metcalfe K, Moore C, Newbury-Ecob R, Omer S, Ouwehand WH, Park S-M, Parker MJ, Pickardt T, Pollard MO, Robert L, Roberts DJ, Sambrook J, Setchfield K, Stiller B, Thornborough C, Toka O, Watkins H, Williams D, Wright M, Mital S, Daubeney PE, Keavney B, Goodship J, Consortium U, Abu-Sulaiman R, Klaassen S, Wright CF, Firth HV, Barrett JC, Devriendt K, FitzPatrick DR, Brook DJ, Study D, Hurles ME. 2016. Distinct genetic architectures for syndromic and nonsyndromic congenital heart defects identified by exome sequencing. Nat Genet 48:1060–1065. doi:10.1038/ng.3627

Silva AP, Ryan DP, Galanty Y, Low JK, Vandevenne M, Jackson SP, Mackay JP. 2016. The N-terminal Region of Chromodomain Helicase DNA-binding Protein 4 (CHD4) Is Essential for Activity and Contains a High Mobility Group (HMG) Box-like-domain That Can Bind Poly(ADP-ribose). J Biol Chem 291:924–938. doi:10.1074/jbc.m115.683227

Sims JK, Wade PA. 2011. Mi-2/NuRD complex function is required for normal S phase progression and assembly of pericentric heterochromatin. Mol Biol Cell 22:3094–3102. doi:10.1091/mbc.e11-03-0258

Sims RJ, Chen C-F, Santos-Rosa H, Kouzarides T, Patel SS, Reinberg D. 2005. Human but Not Yeast CHD1 Binds Directly and Selectively to Histone H3 Methylated at Lysine 4 via Its Tandem Chromodomains. J Biol Chem 280:41789–41792. doi:10.1074/jbc.c500395200

Skene PJ, Hernandez AE, Groudine M, Henikoff S. 2014. The nucleosomal barrier to promoter escape by RNA polymerase II is overcome by the chromatin remodeler Chd1. Elife 3:e02042. doi:10.7554/elife.02042

Smeenk G, Wiegant WW, Vrolijk H, Solari AP, Pastink A, van Attikum H. 2010. The NuRD chromatin–remodeling complex regulates signaling and repair of DNA damage. J Cell Biology 190:741–749. doi:10.1083/jcb.201001048

Sundaramoorthy R, Hughes AL, El-Mkami H, Norman DG, Ferreira H, Owen-Hughes T. 2018. Structure of the chromatin remodelling enzyme Chd1 bound to a ubiquitinylated nucleosome. Elife 7:e35720. doi:10.7554/elife.35720

Sundaramoorthy R, Hughes AL, Singh V, Wiechens N, Ryan DP, El-Mkami H, Petoukhov M, Svergun DI, Treutlein B, Quack S, Fischer M, Michaelis J, Böttcher B, Norman DG, Owen-Hughes T. 2017. Structural reorganization of the chromatin remodeling enzyme Chd1 upon engagement with nucleosomes. Elife 6:e22510. doi:10.7554/elife.22510

Tegunov D, Cramer P. 2018. Real-time cryo-EM data pre-processing with Warp. Biorxiv 338558. doi:10.1101/338558

Tong JK, Hassig CA, Schnitzler GR, Kingston RE, Schreiber SL. 1998. Chromatin deacetylation by an ATP-dependent nucleosome remodelling complex. Nature 395:917–921. doi:10.1038/27699

Tsurusaki Y, Okamoto N, Ohashi H, Kosho T, Imai Y, Hibi-Ko Y, Kaname T, Naritomi K, Kawame H, Wakui K, Fukushima Y, Homma T, Kato M, Hiraki Y, Yamagata T, Yano S, Mizuno S, Sakazume S, Ishii T, Nagai T, Shiina M, Ogata K, Ohta T, Niikawa N, Miyatake S, Okada I, Mizuguchi T, Doi H, Saitsu H, Miyake N, Matsumoto N. 2012. Mutations affecting components of the SWI/SNF complex cause Coffin-Siris syndrome. Nat Genet 44:376. doi:10.1038/ng.2219

Vasudevan D, Chua E, Davey CA. 2010. Crystal Structures of Nucleosome Core Particles Containing the ‘601’ Strong Positioning Sequence. J Mol Biol 403:1–10. doi:10.1016/j.jmb.2010.08.039

Waterhouse AM, Procter JB, Martin DM, Clamp M, Barton GJ. 2009. Jalview Version 2—a multiple sequence alignment editor and analysis workbench. Bioinformatics 25:1189–1191. doi:10.1093/bioinformatics/btp033

Weiss K, Terhal PA, Cohen L, Bruccoleri M, Irving M, Martinez AF, Rosenfeld JA, Machol K, Yang Y, Liu P, Walkiewicz M, Beuten J, Gomez-Ospina N, Haude K, Fong C-T, Enns GM, Bernstein JA, Fan J, Gotway G, Ghorbani M, udy, van Gassen K, Monroe GR, van Haaften G, Basel-Vanagaite L, Yang X-J, Campeau PM, Muenke M. 2016. De Novo Mutations in CHD4, an ATP-Dependent Chromatin Remodeler Gene, Cause an Intellectual Disability Syndrome with Distinctive Dysmorphisms. Am J Hum Genetics 99:934–941. doi:10.1016/j.ajhg.2016.08.001

Willhoft O, Ghoneim M, Lin C-L, Chua EY, Wilkinson M, Chaban Y, Ayala R, McCormack EA, Ocloo L, Rueda DS, Wigley DB. 2018. Structure and dynamics of the yeast SWR1-nucleosome complex. Science 362:eaat7716. doi:10.1126/science.aat7716

Winger J, Nodelman IM, Levendosky RF, Bowman GD. 2018. A twist defect mechanism for ATP-dependent translocation of nucleosomal DNA. Elife 7:e34100. doi:10.7554/elife.34100

Woodage T, Basrai MA, Baxevanis AD, Hieter P, Collins FS. 1997. Characterization of the CHD family of proteins. Proc National Acad Sci 94:11472–11477. doi:10.1073/pnas.94.21.11472

Xia L, Huang W, Bellani M, Seidman MM, Wu K, Fan D, Nie Y, Cai Y, Zhang YW, Yu L-R, Li H, Zahnow CA, Xie W, Yen R-W, Rassool FV, Baylin SB. 2017. CHD4 Has Oncogenic Functions in Initiating and Maintaining Epigenetic Suppression of Multiple Tumor Suppressor Genes. Cancer Cell 31:653-668.e7. doi:10.1016/j.ccell.2017.04.005

Xue Y, Wong J, Moreno GT, Young MK, Côté J, Wang W. 1998. NURD, a Novel Complex with Both ATP-Dependent Chromatin-Remodeling and Histone Deacetylase Activities. Mol Cell 2:851–861. doi:10.1016/s1097-2765(00)80299-3

Yan L, Wang L, Tian Y, Xia X, Chen Z. 2016. Structure and regulation of the chromatin remodeller ISWI. Nature 540:466. doi:10.1038/nature20590

Yan L, Wu H, Li X, Gao N, Chen Z. 2019. Structures of the ISWI–nucleosome complex reveal a conserved mechanism of chromatin remodeling. Nat Struct Mol Biol 1–9. doi:10.1038/s41594-019-0199-9

Zhang Y, LeRoy G, Seelig H-P, Lane WS, Reinberg D. 1998. The Dermatomyositis-Specific Autoantigen Mi2 Is a Component of a Complex Containing Histone Deacetylase and Nucleosome Remodeling Activities. Cell 95:279–289. doi:10.1016/s0092-8674(00)81758-4

Zivanov J, Nakane T, Forsberg BO, Kimanius D, Hagen WJ, Lindahl E, Scheres SH. 2018. New tools for automated high-resolution cryo-EM structure determination in RELION-3. Elife 7:e42166. doi:10.7554/elife.42166

